# Preserving Hidden Hierarchical Structure: Poincaré Distance for Enhanced Genomic Sequence Analysis

**DOI:** 10.1101/2024.10.11.617848

**Authors:** Sarwan Ali, Haris Mansoor, Prakash Chourasia, Imdad Ulla Khan, Murray Pattersn

## Abstract

The analysis of large volumes of molecular (genomic, proteomic, etc.) sequences has become a significant research field, especially after the recent coronavirus pandemic. Although it has proven beneficial to sequence analysis, machine learning (ML) is not without its difficulties, particularly when the feature space becomes highly dimensional. While most ML models operate with the conventional Euclidean distance, the hidden hierarchical structure present among a set of phylogenetically related sequences is difficult to represent in Euclidean space without losing much information or requiring many dimensions. Since such hierarchical structure can be informative to analysis tasks such as clustering and classification, we propose two measures for generating a distance matrix from a set of sequences based on distance in the Poincaré disk model of hyperbolic geometry, or the *Poincaré distance*, for short. Such a distance measure can allow to embedding of even a fully resolved phylogenetic tree in just two dimensions with minimal distortion to any hierarchical structure. Our first approach is based purely on the classical Poincaré distance, while the other approach modifies this distance by combining the Euclidean norms and the dot product between the sequence representations. A thorough analysis of both measures demonstrates its superiority in a variety of genomic and proteomic sequence classification tasks in terms of efficiency, accuracy, predictive performance, and the capacity to capture significant sequence correlations. These approaches perform better than existing state-of-the-art methods across the majority of evaluation metrics.

## 1 Introduction

In the field of bioinformatics, the analysis of molecular (genomic, proteomic, etc.) sequences is a fundamental task that plays a crucial role in understanding their structure, function, and evolution [1,2]. On the other hand, the advancements in machine learning and artificial intelligence has led to the emergence of learning representations for symbolic data [3], such as text [4], graphs [5], and multi-relational data [6]. The use of machine learning (ML) models has become popular due to their ease of use, generalizability, and efficient performance [7,8,9,10,11]. However, since most of the ML models work with Euclidean distance in general, it is difficult to capture hierarchical information such as phylogenetic relationships simply with pairwise Euclidean distances between molecular sequence representations.

Recent methods for designing embeddings for molecular sequences include one-hot encoding (OHE) [12] and *k*-mers spectrum representations [13]. OHE embeddings are highly-dimensional and lack order information [14], while the *k*-mers spectrum representation preserves sequence order but overlooks structural details [15]. Euclidean distance in either the OHE or the *k*-mers spectrum representation fails to consider hierarchical information, impacting downstream classification tasks. Innovative approaches, like kernel matrices, attempt to address this by projecting data into a high-dimensional space [16], yet might not fully capture all nonlinear relationships [16]. Thus, there remains a need for more effective methods that comprehensively capture sequence similarity and other relationships between sequences. To overcome these drawbacks and enhance sequence analysis potential, this research proposes a new distance measure based on distance in the Poincaré disk model of hyperbolic geometry, or the *Poincaré distance*, for short [17]. Hyperbolic geometry has the unique property of preserving both visible and hidden hierarchical structure in the data. By leveraging this property, the Poincaré distance aims to provide a more accurate representation of genomic sequence similarity compared to the conventional Euclidean distance, capturing the complex relationships and dependencies present in the data.

In this study, we compute a kernel matrix for molecular sequences using the Poincaré distance, which captures nonlinear, hierarchical relationships and is more robust to outliers compared to traditional dot product-based kernel matrices.

Furthermore, we propose a modified version of the Poincaré distance, denoted as M-Poincaré, which combines Euclidean norms and the dot product between the molecular sequence representations. This modification aims to enhance the representation of sequence similarities by incorporating both hyperbolic and Euclidean geometry. This strikes a balance between the simplicity of computing the representation and learning from it, with the ability to capture a more complex (hierarchical) structure and what can be learned from this. The suggested Poincaré and M-Poincaré distances hold great promise and have real-world applications in disciplines such as medicine, biotechnology, and information retrieval. Offering better genomic sequence similarity representation and enhancing tasks like sequence classification, functional annotation, and evolutionary analysis.

Overall, we propose a novel distance-based kernel matrix using Poincaré distance, its evaluation and validation on a diverse set of genomic and proteomic sequence data sets, and the practical implications in real-world applications. In summary, the contributions are as follows:

1. **Novel Distance Measure:** This paper proposes two novel distance measures based on the Poincaré disk model of hyperbolic geometry for molecular sequence analysis. Unlike conventional Euclidean distance, the proposed distance measure takes advantage of the unique properties of the hyperbolic space to capture the complex relationships and dependencies within molecular sequence data. By utilizing hyperbolic geometry, which is known for its ability to represent hierarchical (tree-like) structure, the proposed distance measure offers a more accurate representation of similarity for sets of phylogenetically related sequences.
2. **Kernel Matrix Computation:** Using the Poincaré distance matrix, we design a kernel matrix using the Gaussian kernel, which shows superior performance compared to the *k*-mers spectrum-based kernel computation method from the literature.
3. **Evaluation and Validation:** This paper extensively evaluates the effectiveness of the proposed distance measures on three sets of genomic (or proteomic) sequences sequences to cover a broad range of analysis cases. We compare the results for the Poincaré kernel and M-Poincaré kernel with several recently proposed baselines and show the superiority of the proposed distance measures in terms of accuracy, efficiency, and the ability to capture meaningful sequence relationships.
4. **Real-World Applications:** The contributions of this paper extends beyond theoretical advancements and has practical implications in real-world applications. Accurate sequence (in the general sense, *i*.*e*., beyond just molecular sequences) comparison is crucial in various domains, such as genomics, evolutionary biology, and recommendation systems.

The rest of the paper is organized as follows: Section 2 contains the literature review for the proposed method. This paper’s contribution is discussed in Section 3. The dataset statistics and experimental setup detail is given in Section 4. The results for proposed and baseline models are reported in Section 5. Finally, the conclusion of this study is given in Section 6.

## 2 Related Work

Traditional distance measures, such as Euclidean distance [18], Hamming distance [19], and cosine similarity [20], have been widely used in molecular sequence analysis. These measures are based on the assumption of sequences being represented as vectors in an Euclidean space [21]. While they have proven to be effective in certain applications, they suffer from limitations for sequences with complex structures and dependencies [21]. For example, Euclidean distance fails to capture the hierarchical relationships and tree-like structures that are prevalent in many sequence datasets in a natural way, requiring many more dimensions to capture it. Moreover, these measures often struggle to handle high-dimensional sequence data and may produce suboptimal results [22].

Recent research has explored the use of hyperbolic geometry in sequence analysis [21,17]. Hyperbolic space offers unique properties that make it well-suited for capturing hierarchical and tree-like structures [21]. Several studies have demonstrated the effectiveness of hyperbolic representations in tasks such as text classification [23], network analysis [24], and knowledge graph embeddings [25]. Graph-based approaches in sequence analysis capture complex relationships by representing sequences as nodes, with edges [26] indicating similarity (k-nearest neighbor or kNN graphs) or interactions [27], such as protein-protein binding. Despite their potential, these methods can be computationally intensive and may struggle with large-scale datasets and capturing hierarchical information efficiently. Embedding-based methods project sequences into low-dimensional vector spaces, reflecting similarity between molecular sequences [28,29,30,31,32,33,34,35,36]. Techniques like *k*-mers spectrum [14,13] have shown reasonable performance in various tasks but can be computationally expensive and may fail to preserve hierarchical and structural relationships effectively.

A common approach to computing sequence similarity is generating a kernel or gram matrix [14,16]. This can be computationally expensive, but some methods improve efficiency by using the dot product of sequence spectra [16]. These matrices [14] are used in classifiers like SVMs or through kernel PCA [37]. However, they often miss hierarchical and structural information in molecular sequences. Models like Protein Bert [38] and SeqVec [39] aim to address this for proteins but struggle to generalize across different functions and perform poorly on nucleotide sequences due to varying biological properties.

## 3 Proposed Approach

Given an abstract space 𝒳 (e.g., proteins, etc.), function : 𝒳 × 𝒳 → ℝ is called a kernel function. Kernel functions are used to quantify the similarity between a pair of objects *x, y* ∈ 𝒳. We propose a kernel function using two distance methods, namely Poincaré distance and Modified Poincaré (or M-Poincaré) distance in hyperbolic space. Hyperbolic geometry is a non-Euclidean geometry that studies spaces of constant negative curvature. Geodesics and distances are also curved in hyperbolic geometry, which are generalizations of straight lines in Euclidean geometry Figure 1 (b). The underlying hyperbolic geometry allows us to learn similarity better than Euclidean space. Figure 1 (a) shows a visualization of a particular type of tiling (isosceles) in hyperbolic geometry. In hyperbolic isosceles tiling, the tiles used are hyperbolic isosceles triangles: they have two equal sides and angles. These triangles are created by cutting a hyperbolic plane into congruent triangles, each having two sides of the same length.

**Fig 1:**
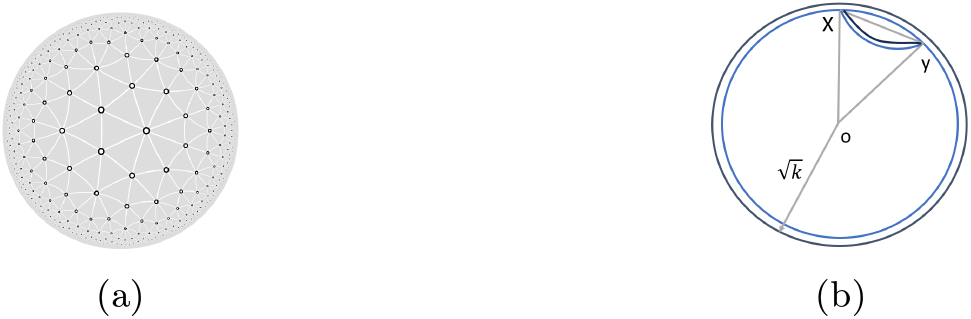
(a) Hyperboloid isosceles tiling, each triangle has two equal sides and equal angles, we can observe negative curvature and exponential growth. (b) As the curvature (−1*/K*) of the Poincare disk decreases, the distance between points *x* and *y* connected by geodesics (shortest paths) on the disk increases.

We divide our method into three steps, where we first compute the pairwise distance between the data points. The second step involves computing the kernel values using the Gaussian kernel. In the end, we use kernel PCA to get the final embeddings, which are then used for underlying supervised analysis.

### 3.1 Distance Computation

We will now describe both of these distance methods in detail. The Poincaré distance *d*(*x, y*) between a pair *x, y* of vectors is defined as follows:

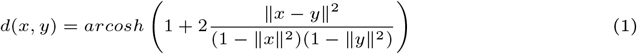

The notation ∥· ∥ denotes the Euclidean norm, which is the square root of the sum of squared differences between corresponding elements, arcosh(*z*) is the inverse hyperbolic cosine function, a.k.a the area hyperbolic cosine function. It is the inverse function of the hyperbolic cosine function cosh(*z*) and returns the value whose hyperbolic cosine is *z*. Therefore, the Poincaré distance between vectors *x* and *y* is calculated by taking the Euclidean distance between them, scaling it using the terms (1 − ∥*x*∥^2^)(1 − ∥*y*∥^2^), multiplying by 2, adding 1, and then taking the inverse hyperbolic cosine of the result.

We propose a distance function (a modified form of Poincaré distance) that combines elements of Euclidean norms and dot products. Our distance function calculates the distance between two vectors *x* and *y*. It is a measure of dissimilarity, defined as:

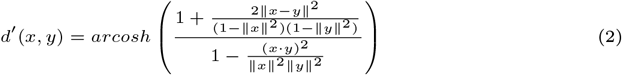

where *d*′(*x, y*) represents the modified Poincaré distance between vectors *x* and *y*, and *x* · *y* represents the dot product of vectors *x* and *y*.

The difference of modified Poincaré distance formula *d*′ in Equation 2 from the original Poincaré distance formula *d* in Equation 1 is in the denominator of the fraction. In the standard Poincaré distance formula, the denominator is defined as the product of the norms of the vectors x and y. In the modified formula (Equation 2), the denominator is defined as the dot product of the vectors x and y divided by the product of their norms. This modification in the denominator allows for the inclusion of the dot product term, which can capture additional information about the similarity or dissimilarity between the vectors x and y based on their orientations. Hence, in addition to magnitude, it incorporates the orientation information of the vectors.

The M-Poincaré distance represents a significant advancement in sequence analysis by seamlessly integrating the strengths of hyperbolic and Euclidean geometries. By combining hyperbolic geometry’s hierarchical structure preservation with Euclidean geometry’s compatibility with high-dimensional data, M-Poincaré offers a comprehensive representation of sequence relationships. It unites hyperbolic divergence and Euclidean norms, effectively capturing both global and local features in molecular sequences, addressing the limitations of methods relying on a single geometry. This innovative hybrid approach provides a nuanced perspective on evolutionary dynamics, making M-Poincaré a superior choice for sequence analysis that bridges existing gaps in current methodologies.

### 3.2 Theoretical Properties

A kernel function typically satisfies the following two properties: non-negativity, and symmetry. A kernel with these properties will loosely have the interpretation as a similarity quantification. Both *d*(*x, y*) and *d*′(*x, y*) represent a similarity metric on input space.

- Non-Negativity: *d*(*x, y*) *>* 0, *d*(*x, y*)′ *>* 0
- Symmetry: *d*(*x, y*) = *d*(*y, x*), *d*′(*x, y*) = *d*′(*y, x*)

*Remark 1*. Note that the difference between *d*(*x, y*) and *d*′(*x, y*) is that the modified Poincaré distance has an inner product in the denominator which accounts for the extra cosine similarity measure.

#### Input Embedding Representation

In this study, we utilize the *k*-mers spectrum [14] as the input to the Poincaré and modified Poincaré distance functions. The *k*-mers spectrum is a representation of sequences that captures the frequency of occurrence of all possible subsequences of length k, where *k >* 0. Formally, given an alphabet *Σ* (unique amino acid characters), the length of the *k*-mers spectrum is equal to |*Σ*|^*k*^ (where *Σ* = 20 for protein sequences and *Σ* = 4 for DNA sequences.). The *k*-mers spectrum provides a compact representation of sequences by encoding the information about the frequencies of all possible k-mers. Offers several advantages over one-hot encoding (OHE) [12], which suffers from a high-dimensional and sparse representation, especially for large alphabets or long sequences. The *k*-mers spectrum also retains important positional and contextual information within the sequence. Unlike one-hot encoding, which treats each element independently (hence does not preserve the order of amino acids/nucleotides within molecular sequences), the *k*-mers spectrum considers the occurrence of subsequences and their frequencies. This enables the detection of patterns, motifs, and dependencies within the sequence, which are crucial for various sequence analysis tasks.

*Remark 2*. We also computed results using the one-hot encoding-based representation of molecular sequences as input. However, the results were not encouraging Title Suppressed Due to Excessive Length 7 compared to the *k*-mers-based spectrum (due to the reasons discussed above) for the proposed method. Hence we did not include those results in this paper.

### 3.3 Kernel Matrix From Poincaré Distance

Since Equation 1 and 2 are used to design the distance matrix between each pair of sequences, we cannot use them as kernel matrix. Generating a kernel matrix from the Poincaré-based distance matrix offers technical justifications such as capturing nonlinear relationships, flexibility with different ML algorithms, preserving essential information, and potential performance enhancements [17]. Detailed benefits of designing kernel matrix from the Poincaré distance function are given in Section 3.6. To construct the kernel matrix based on the Poincaré distance or M-Poincaré distance, we employ a Gaussian kernel. The Gaussian kernel, also known as the Radial Basis Function (RBF) kernel [40], is a popular choice for capturing complex relationships and nonlinear patterns in data. These kernel matrices are then used to design the embeddings using kernel PCA [37].

The kernel matrix is constructed by applying the Gaussian kernel function to the pairwise distances between sequences using either the Poincaré distance or the M-Poincaré distance. The Gaussian kernel is defined as:

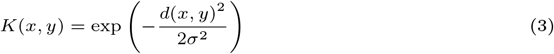

where *d*(*x, y*) represents either the Poincaré distance or the M-Poincaré distance between vectors *x* and *y*, and *σ* is a user-defined width of the kernel.

The Gaussian kernel assigns a similarity value between 0 and 1 to each pair of sequences based on their distance. The closer the sequences are, the higher the similarity value. The exponential term in Equation 3 ensures that the similarity decays exponentially with increasing distance. By applying the Gaussian kernel to the pairwise distances, we obtain a symmetric positive semi-definite kernel matrix. This matrix captures the similarity structure among the sequences, with higher values indicating similarity and lower values indicating dissimilarity. The pseudocode for computing the Poincaré kernel is given in Algorithm 1 and Figure 2 shows the flow chart. The algorithm takes a set of molecular sequences as input and computes *k*-mers spectrum-based numerical embeddings (line 1). It then iterates for all pairs of embeddings (upper triangle only due to symmetry property), computes the Poincaré distance using Equation 1 (line 8), and applies Gaussian kernel to get the final kernel value (line 9). The same process is performed for M-Poincaré by replacing Equation 1 (line 8) with Equation 2.

#### Algorithm 1

Poincaré Kernel Matrix

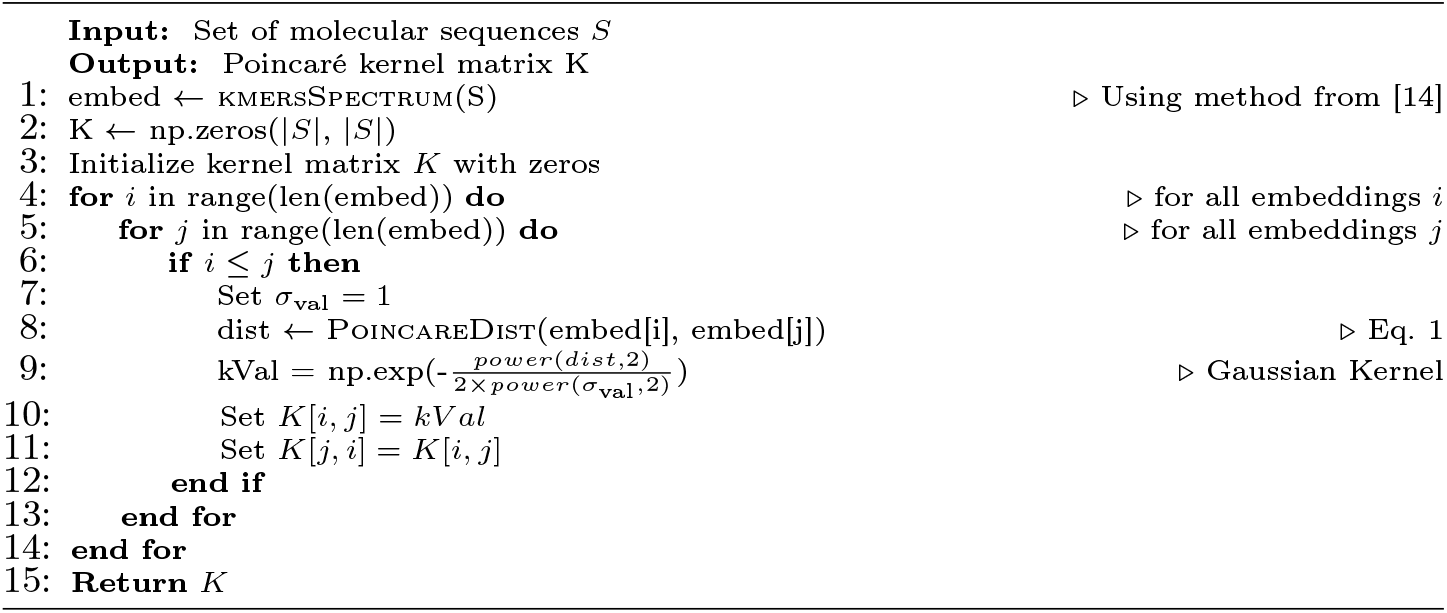

**Fig 2:**
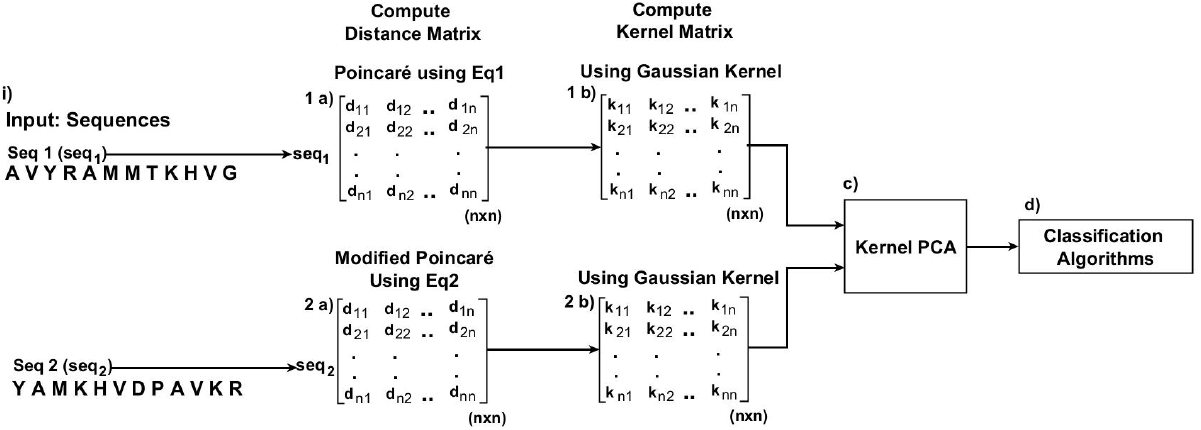
Poincaré and Modified Poincaré Distance-based classification Workflow.

### 3.4 Kernel PCA [37]

Kernel PCA creates low-dimensional embeddings of sequences by leveraging the kernel matrix generated from distances like Poincaré or M-Poincaré. It extends PCA by utilizing the kernel matrix’s eigenvectors to capture nonlinear patterns and structures. By applying kernel PCA, hierarchical or tree-like structures within data and complex nonlinear patterns can be accurately captured and embedded. Incorporating hyperbolic geometry in distance calculations and the Gaussian kernel enhances the expressiveness and informativeness of these embeddings, benefiting tasks such as clustering and classification. A detailed justification of using Kernel Matrix is given in Section 3.6. Further detailed discussion on how the Poincaré-based distance preserves hierarchical information is given in Section 3.5.

### 3.5 Discussion On How the Poincaré-based distance preserves hierarchical information

The utilization of the Poincaré-based distance, rooted in hyperbolic geometry, provides a distinctive perspective for capturing hierarchical connections within molecular sequences. This preservation of hierarchy stands out especially when compared to the traditional Euclidean distance.

Hyperbolic Geometry’s Hierarchical Preservation: Hyperbolic geometry functions within a space that deviates from Euclidean norms by not adhering to the parallel postulate. This departure from Euclidean principles grants hyperbolic space unique qualities that are particularly adept at capturing hierarchies. Notably, volume increases exponentially in hyperbolic space as one moves away from a given point, resulting in a “concentration effect.” This mirrors the way molecular sequences evolve, with new species or variations branching out on the evolutionary tree. The Poincaré distance leverages this concentration effect, making it a powerful tool for representing evolutionary hierarchies. As sequences diverge over time, the Poincaré distance amplifies their differences, effectively representing their hierarchical relationships.

Limitations of Euclidean Distance in Capturing Hierarchy: In contrast, the Euclidean distance operates in a flat Euclidean space, where distances grow linearly which imposes limitations on capturing hierarchies. For instance, as species evolve and diverge, the linear distance growth in Euclidean space may inadequately capture the extent of divergence. This results in a loss of hierarchical information, as linear scaling does not align with the exponential branching characteristic of biological evolution. This leads to a struggle to encapsulate the intricate branching and divergence inherent in molecular sequences.

#### Illustrative Scenario

Imagine species A, B, and C evolving from a common ancestor. In hyperbolic space, the Poincaré distance between A and B, and between B and C, will be noticeably smaller compared to the combined distances between A and C in a pairwise Euclidean comparison. This mirrors the hierarchical relationship, indicating that A and B share a more recent common ancestor than A and C. The Poincaré-based distance inherently preserves this evolutionary tree structure, accurately reflecting the dynamics of hierarchy.

### 3.6 Justification of Using Kernel Matrix

Generating a kernel matrix from the Poincaré-based distance matrix offers several technical justifications and benefits:

1. **Nonlinearity:** The Poincaré distance captures the underlying hyperbolic geometry, which is a non-Euclidean space. By constructing a kernel matrix based on the Poincaré distance, we can implicitly map the data into a high-dimensional feature space, allowing for the capture of complex nonlinear relationships. This is especially beneficial when dealing with hierarchical or tree-like structures, which can be present in molecular sequences.
2. **Capturing Complex Relationships:** The Gaussian kernel used in generating the kernel matrix allows for the capture of complex relationships between sequences. It assigns higher similarity values to similar sequences and lower values to dissimilar ones. This enables the representation of intricate patterns and structures in the data, which may not be captured by linear methods.
3. **Theoretical Properties of Gaussian Kernel:** Since we are using the popular Gaussian kernel, all the underlying theoretical properties including (1) Reproducing Kernel Hilbert Space “RKHS” [41] (allows the use of kernel methods, such as support vector machines, for efficient and effective learning from data), (2) Universal Approximation [42] (can approximate any continuous function arbitrarily well in the RKHS, making it a powerful tool for modeling complex relationships and capturing nonlinear patterns in data), (3) Mercer’s Theorem [43] (guarantees that the kernel matrix is positive semi-definite), (4) Smoothness and Continuity [44] (assigns higher similarity values to points that are closer together and lower values to points that are farther apart, allowing the Gaussian kernel to capture local relationships and gradually decrease the influence of distant data points), and (5) Sensitivity to Variations [45] (capture fine-grained changes in the data distribution and assign different similarity values accordingly).
4. **Flexibility and Generalization:** The kernel matrix derived from the Poincaré-based distance matrix can be used with various machine learning algorithms that operate on kernel matrices. This provides flexibility and allows for the application of a wide range of techniques, such as kernel PCA and kernel SVM, etc. By utilizing these algorithms, we can leverage the expressive power of the kernel matrix to address different tasks, including dimensionality reduction and classification.
5. **Preserving Nonlinear Information:** Kernel PCA, performed on the kernel matrix, captures the essential nonlinear information in the data. By projecting the data onto the principal components, we retain the discriminative information while reducing the dimensionality. This can be useful when visualizing the data or when dealing with high-dimensional datasets.

## 4 Experimental Evaluation

This section describes the experimental setup, dataset used, and evaluation metrics. All experiments are conducted using Python on a system equipped with a Core i5 processor running at 2.4 GHz, 32 GB of memory, and the Windows 10 OS.

### Dataset Statistics

We use 3 real-world molecular sequence data comprised of both proteins and nucleotide sequences as follows:

#### Spike7k

The spike protein sequences of the SARS-CoV-2 virus total 7000 sequences are distributed among a total of 22 coronavirus Lineage [46].

#### Human DNA

A total of 4380 Unaligned nucleotide sequences to classify among 7 gene families to which humans belong [47].

#### Coronavirus Host

The spike protein sequences belonging to various clades of the Coronaviridae family are accompanied by the infected host label total of 21 labels e.g. Humans, Bats, Chickens, etc. A total of 5558 sequences are includedViPR [48], GISAID [46].

### Baseline Methods

We carefully selected several well-known baselines from the literature comprised of different embedding generation categories including feature engineering (i.e. PWM2Vec [28]), traditional kernel matrix generation with kernel PCA (i.e. String Kernel [16]), neural networks (i.e. WDGRL [30]), pre-trained language model (i.e. SeqVec [39]), and pre-trained transformer (i.e. Protein BERT [38]) for protein sequences.

#### Evaluation Metrics

To evaluate different models, we use average accuracy, precision, recall, F1 (weighted), F1 (macro), Receiver Operator Characteristic Curve (ROC) Area Under the Curve (AUC), and training runtime. For the metrics designed for the binary classification problem, we use the one-vs-rest approach for multi-class classification. Supervised analysis is performed using different linear and non-linear classifiers including Support Vector Machine (SVM), Naive Bayes (NB), Multi-Layer Perceptron (MLP), K-Nearest Neighbors (KNN), Random Forest (RF), Logistic Regression (LR), and Decision Tree (DT).

## 5 Results And Discussion

The classification results (averaged over 5 runs) for the proposed and baseline methods are given in Table 1 for Spike7k data. We can observe that the proposed Poincaré and M-Poincaré based methods consistently outperform the baseline models in terms of predictive performance. Across most evaluation metrics, these methods achieve higher scores compared to other embedding methods.

**Table 1:**
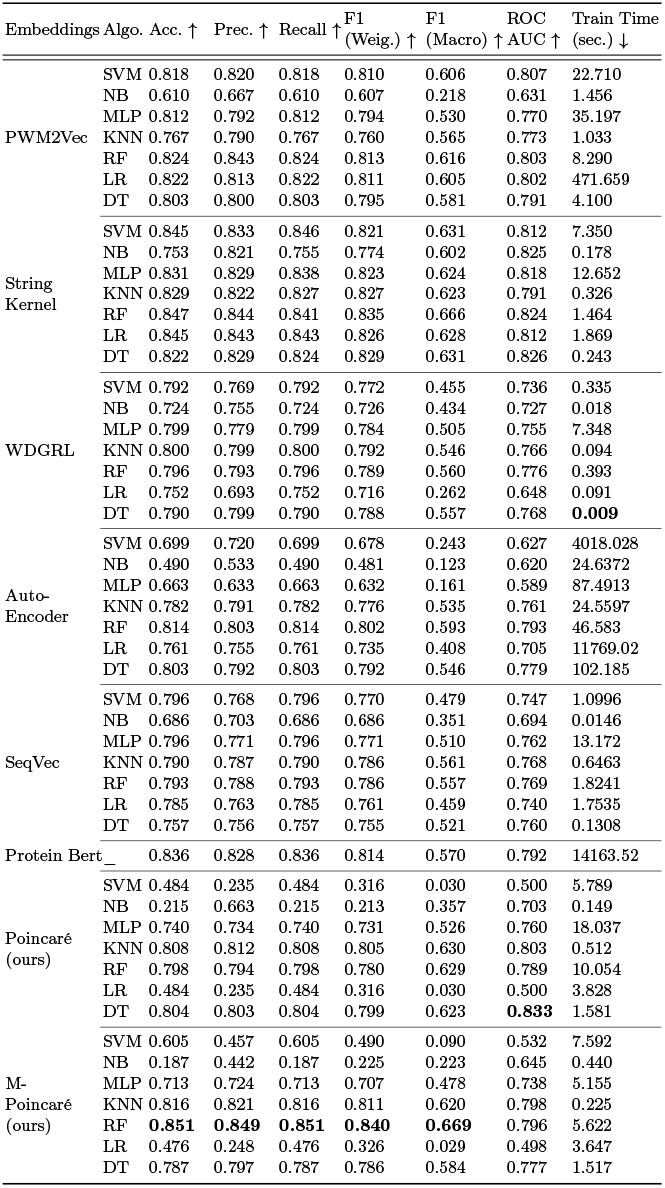
Classification results (averaged over 5 runs) on **Spike7k** dataset for different evaluation metrics. The best values are shown in bold.

The average classification results for the Human DNA (nucleotides) dataset are reported in Table 2. Here we can observe that the proposed Poincaré kernel outperforms all other methods in terms of average accuracy, recall, weighted F1, macro F1, and ROC-AUC. For precision, the M-Poincaré performs the best. An important behavior to note here is that the pretrained Protein Bert performed very poorly on this dataset compared to its performance on Spike7k and coronavirus host datasets. The reason behind this behavior is the fact that the Protein Bert model is specifically trained on the protein sequence data. Since Human DNA consists of nucleotide sequences, this method failed to generalize in this case. On the other hand, the proposed Poincaré kernel and M-Poincaré kernel-based methods show the highest performance compared to the baselines.

**Table 2:**
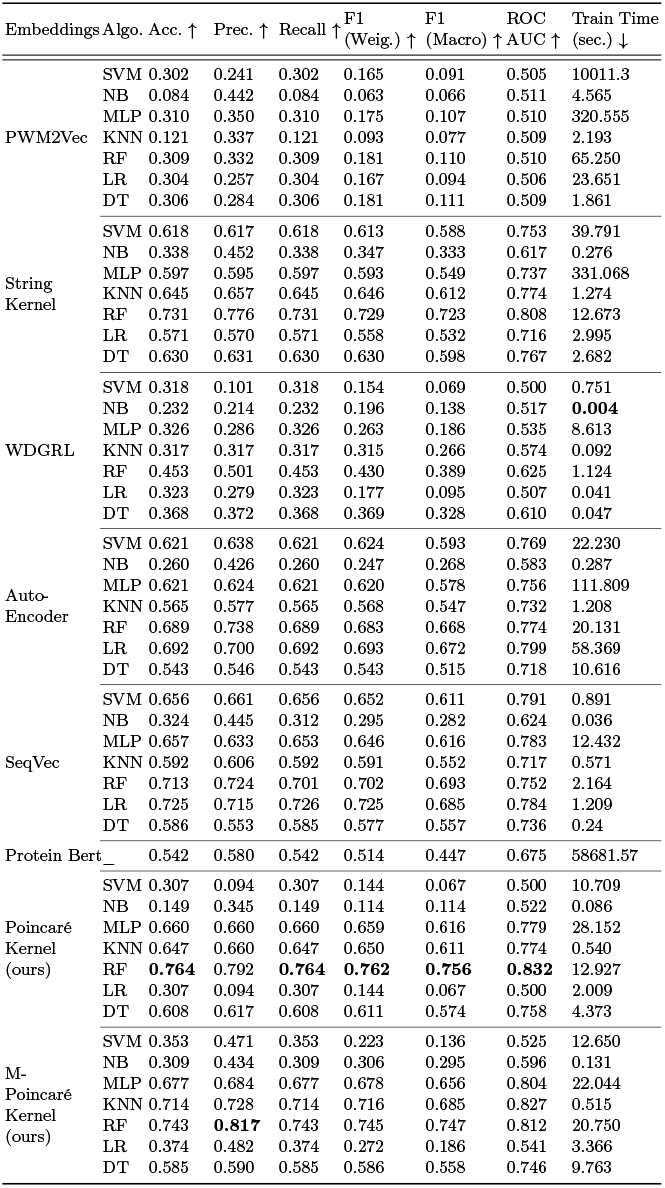
Classification results (averaged over 5 runs) on **Human DNA** dataset for different evaluation metrics. The best values are shown in bold.

The average classification results for the coronavirus host (spike proteins) dataset are reported in Table 3. Here we can observe that the proposed Poincaré kernel outperforms all other methods in terms of average accuracy, precision, recall, weighted F1, and ROC-AUC. For macro F1, the baseline PWM2Vec method performs the best. Overall the proposed Poincaré kernel and M-Poincaré kernel outperform all baselines for both proteins and nucleotide datasets in terms of average classification results. We also compared the results for Euclidean vs. Hyperbolic embeddings (Poincaré and M-Poincaré) in Section 5.1.

**Table 3:**
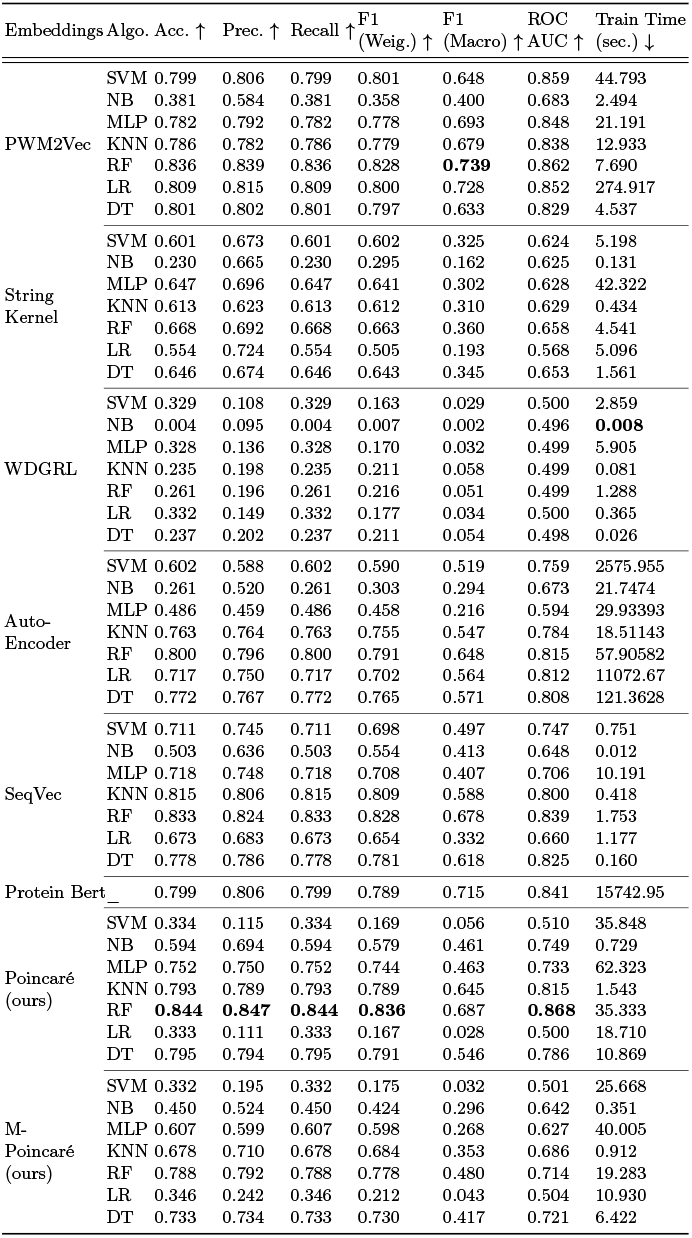
Classification results (averaged over 5 runs) for different evaluation metrics for **Coronavirus Host Dataset**. The best values are in bold.

### 5.1 Euclidean vs. Hyperbolic Embedding Results

To analyze if the proposed embedding method from hyperbolic geometry (i.e. Poincaré and M-Poincaré), we generate the embeddings using Euclidean (i.e. L2) distance between the input vectors as an alternative to the Poincaré and M-Poincaré distances from Equation (1) and Equation (2), respectively, from the main paper. The remaining steps of this Euclidean-distance-based representation remain the same as discussed in Algorithm 1 in the main paper.

The results comparison for Euclidean vs. Hyperbolic representations are reported in Table 4 for Spike7k, Human DNA, and Covid Host dataset. We can observe that for all the datasets, the Poincaré and M-Poincaré-based representations significantly outperform the L2-based Euclidean representations for the supervised analysis. This behavior highlights that there is indeed a hidden hierarchical structure in the molecular sequences, which vanishes when we represent the sequences in Euclidean space. However, those structures are better preserved in the case of hyperbolic geometry.

**Table 4:**
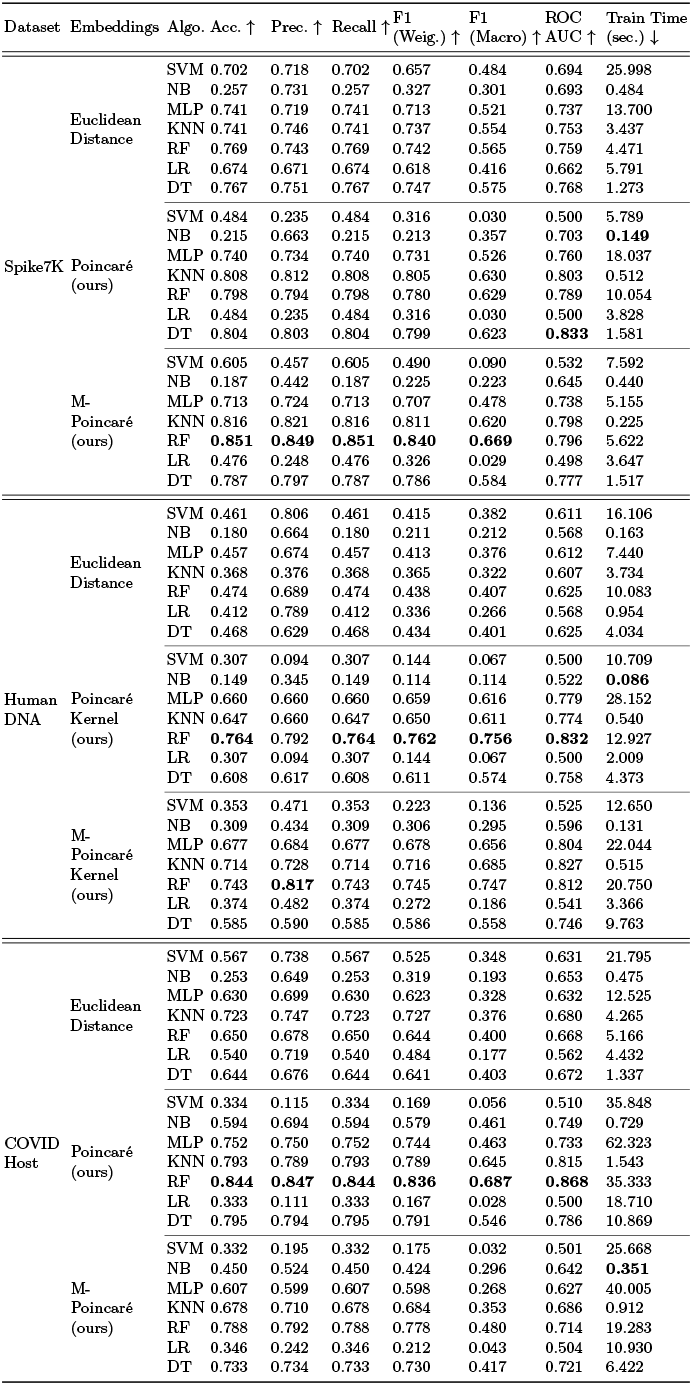
Euclidean vs. Hyperbolic geometry-based embeddings classification results. The best values for respective datasets are shown in bold.

An interesting observation can be observed in the case of the Human DNA dataset (in Table 4), which comprises nucleotide sequences. Since the nucleotide sequences provide direct information about the DNA composition of organisms, they are well-suited for studying evolutionary relationships at the genetic level, including mutations, genetic diversity, and population genetics. Therefore, nucleotide sequences can be used to trace evolutionary events such as gene duplications, insertions, deletions, and recombination. Based on these analyses, preserving the hierarchical information becomes more important when dealing with the nucleotide sequences. However, we can observe in Table 4 that Euclidean distance-based embeddings show significantly lower results compared to hyperbolic distance-based embeddings. This behavior highlights the fact that if a certain type of data contains more (and important) hierarchical information, preserving that information is more important, and ignoring such information can reduce the performance of the generated embeddings.

Compared to the nucleotide sequences, the protein sequences might not capture certain types of genetic changes (e.g., silent mutations) that nucleotide sequences do. Therefore, the hidden hierarchical information in the case of protein sequences might not be as important compared to the nucleotide sequences. Due to these reasons, we can observe in Table 4 that although the Euclidean-distance-based embeddings (for Spike7k, and Coronavirus Host protein sequence datasets) show lower predictive accuracies compared to the hyperbolic distance-based representations, the difference in the results is not too significant compared to the difference observed in the case of Human DNA (nucleotide sequence) dataset.

### 5.2 Statistical Significance

Since the computation of the results involves splitting the data randomly into training and testing sets, the statistical significance of the results is needed. We used the student t-test and observed the *p*-values using averages (reported in the above results table) and standard deviations (SD) of 5 runs. Note that the SD values (of 5 runs with different random splits) for all metrics are very small i.e. *<* 0.002 in the majority of the cases. Therefore, we noted that *p*-values were *<* 0.05 in the majority of the cases (because SD values are very low), confirming the statistical significance of the results.

### 5.3 Inter-Class Embedding Interaction

We use heat maps to analyze further if the proposed kernel can identify different classes better. We took the average of the similarity values to compute a single value for each pair of classes and then computed the pairwise cosine similarity of different class’s embeddings with one another. The heat map is further normalized between [0-1] to the identity pattern. The heatmaps for the *k*-mers spectrum-based embedding [13] and its comparison with the proposed M-Poincaré Kernel-based embeddings are reported in Figure 3, 4 and 5. For the *k*-mers spectrum-based heatmap, the embeddings for different labels have high similarity among them, which means that it is difficult to distinguish between different classes due to high pairwise similarities among their vectors. On the contrary, it is distinguishable for the proposed M-Poincaré Kernel-based embeddings. This essentially means that the embeddings from similar classes are highly similar to each other, indicating that the proposed methodology can accurately identify similar classes and different classes.

**Fig 3:**
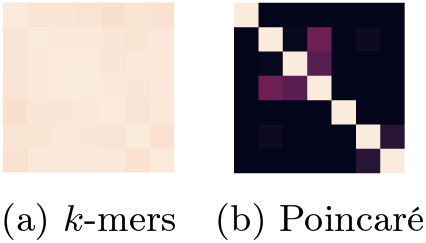
Human DNA.

**Fig 4:**
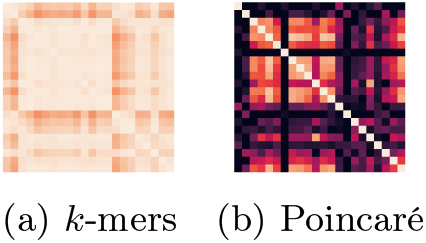
Spike7K.

**Fig 5:**
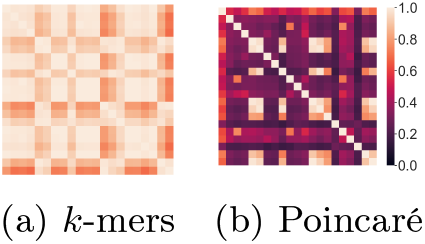
COVID Host. Heatmap for classes in different datasets.

## 6 Conclusion

In this study, we have addressed the limitations of traditional Euclidean-based distance measurements discussed the concept of hyperbolic geometry, and proposed the use of Poincaré distance as a more effective and meaningful measure. By leveraging the unique properties of hyperbolic space, the Poincaré distance preserves the hierarchical structures present in molecular sequences. Furthermore, we introduced a modified version of the Poincaré distance, known as M-Poincaré, which combines Euclidean norms and the dot product between sequence representations. Our proposed distance measure shows superior performance in various evaluation metrics compared to existing methods. The practical implications of our research extend to domains such as medicine, biotechnology, and information retrieval, where accurate sequence comparison is crucial. Future research can explore the application of these methods in other domains along with interpretability studies.

